# Cornelia de Lange Syndrome-associated mutations in Smc1 cause both sister chromatid cohesion and cohesion-independent defects

**DOI:** 10.1101/263418

**Authors:** Jingrong Chen, Frank Wu, Dean Dawson, Susannah Rankin

## Abstract

Cornelia de Lange Syndrome is a pervasive developmental disorder characterized by limb truncations, craniofacial abnormalities, and cognitive delays. This syndrome is a member of a class of developmental disorders referred to as *cohesinopathies*, which result from mutations in the genes encoding subunits or regulators of the cohesin complex. The phenotypic consequences of these mutations may reflect the critical role that cohesin plays in chromosome structure, its ability to tether sister chromatids together during cell cycle progression, or some combination of both. Here we show that a sensitized assay for chromosome loss in budding yeast can be used to assess the impact of Cornelia de Lange syndrome (CdLS)-associated mutations in the core cohesin subunit Smc1 on cohesin function. We find that the CdLS-associated mutations can be grouped into two classes based on their impact on chromosome segregation. One class of mutations includes those that are defective in promoting accurate chromosome segregation, some no better than the null allele. Another class promotes both accurate chromosome cohesion and segregation. Strikingly, the mutations that have no impact chromosome dynamics in this assay are clustered near each other in the context of the folded SMC1 protein suggesting a previously uncharacterized region of functional importance in higher eukaryotes. This analysis illustrates how budding yeast can be used to elucidate mechanisms important in human health and development.

## Introduction

Cornelia de Lange Syndrome, is characterized by limb truncations, craniofacial abnormalities, and, often, severe developmental and cognitive delays. Cornelia de Lange Syndrome (CdLS) was originally attributed to mutations in the NippedB^Scc2^ subunit of the dimeric cohesin loader, Scc2/Scc4 ^1,2^. In recent years, milder variants of CdLS have been attributed to mutations in the gene encoding the cohesin deacetylase, HDAC8, as well as those encoding the structural subunits of the cohesin complex, Smc1 and Smc3 ^3-6^. Smc1 and Smc3 are large coiled-coil containing proteins that interact to form a heterodimer that along with the Rad21 and SA subunits forms the core cohesin complex. Cohesin binds chromatin and tethers sister chromatids together from the time they are made during DNA replication until chromosome segregation at anaphase ^7^. Cohesion between sister chromatids promotes their accurate segregation during cell division, and mediates certain kinds of DNA repair. In recent years, it has become evident that the cohesin complex also has profound effects on gene regulation and thus cellular and developmental phenotypes ^8^. Cohesin is thought to control gene expression by regulating chromosome structure, controlling the long-range interaction between genes and regulatory elements brought together in space by the formation of chromosome loops and topologically associated domains ^9^. Some studies also suggest a direct interaction between cohesin and the transcriptional machinery. How cohesin is controlled to ensure proper regulation of gene expression is not known.

How do abnormalities in the cohesion apparatus lead to developmental disorders? Several possible mechanisms have been suggested ^6,8,10^. First, insufficient cohesion could lead to delays in cell cycle progression due to checkpoint activation. This, in turn, could cause failure to fully populate tissues at appropriate developmental windows, resulting in developmental defects. Second, aberrant cohesion might affect expression of developmental regulators through its impacts on chromosome structure. Third, the mutations in cohesion proteins might affect their direct interaction with the transcription machinery, altering the expression of key developmental regulators. Finally, cohesin defects could affect the integrity of rDNA repeats, rDNA condensation or nucleolar structure in ways that affect protein translation. The phenotypes could also result from some combination of all of these mechanisms. Understanding how defects in cohesion machinery lead to developmental defects is a major question in the cohesinopathy field. Whereas sister chromatid cohesion failure is consistently evident in Robert’s Syndrome ^11^ this is not the case with CdLS. Analysis of patient cells and cells from mouse and *Drosophila* models with mutations in homologs of *NIPBL*, which encodes NippedB^Scc2^, have shown mixed results in assays for sister chromatid cohesion, with one study reporting cohesion loss but other studies reporting no cohesion defects ^12-14^. In contrast, heterozygous mutations in *NIPBL* result in changes to transcriptional programs ^15^.

CdLS-associated mutations in the Smc1 subunit of cohesin, encoded by the *SMC1A* gene (also known as *SMC1L1*) in humans (*CDLS2* [MIM: 300590]), typically encode amino acid substitutions or small in-frame deletions ^3,5,16^. The *SMC1A* gene is on the pseudoautosomal region of the X chromosome, and is thought to escape X inactivation ^17^. Therefore, the phenotypes in female patients likely reflect the expression of both maternal and paternal alleles, while male patients express only the allele on their single X chromosome. The phenotypes observed in female patients could reflect dominant-negative activity of the mutant protein. Intriguingly, some of the CdLC-associated *SMC1A* mutations affect residues that are highly evolutionarily conserved, but have no previously ascribed function. It is possible that these residues represent sites of direct interaction between the cellular machinery that affects CdLS and the cohesion apparatus. Alternatively, these sites might represent previously uncharacterized functional motifs critical to the tethering activity of the cohesin complex.

How do mutations in *SMC1A* result in developmental defects? Not all cohesinopathy-associated mutations have shown cohesion defects in laboratory assays ^14^. The absence of cohesion defects in cells bearing CdLS mutations could suggest that failures in sister chromatid cohesion are not the immediate cause of the disorder. Alternatively, cohesion defects could contribute to the phenotype, but since severe losses in cohesion would be lethal, only mutations with mild cohesion defects could be tolerated, and these mild defects might not be detected in cohesion assays. Finally, it is possible that CdLS-associated mutations in *SMC1A* have no impact on sister chromatid cohesion, but contribute to defects in chromosome structure only, without impacting sister chromatid tethering. To rigorously test whether CdLS-associated mutations in *SMC1A* compromise sister chromatid cohesion we used sensitive assays available in budding yeast to analyze directly the impacts of these mutations on chromosome segregation, DNA repair, and sister chromatid cohesion. The results demonstrate that the ability of cohesin to tether sister chromatids is compromised by many of the CdLS-associated mutations. In some cases, however, the mutations cause no measurable effect even in a sensitized, quantitative assay for chromosome segregation, suggesting the intriguing possibility that the mutations lie in residues that affect interaction of the cohesin complex with factors that influence development independently of sister chromatid cohesion.

## Methods

### Strains and Primers

The genotypes of the strains used in this work are shown in Table S1. All strains are derived from YNN281, with the exception of those used in Figure 4, which are primarily of S288C and W303 ancestry and were derived in the RE Esposito laboratory ^18^.

Primers used to build and confirm the strains are shown in Table S2. Yeast media, culture techniques, and strain construction methods are as described in ^19^.

### Strain construction

Polymerase chain reaction (PCR)-based methods were used to create deletions of open reading frames and mutated versions of *SMC1*. Strains containing mutations in the *SMC1* gene were built by a modified two-step gene replacement method. Briefly, *SMC1* gene fragments were amplified using high-fidelity fusion PCR with primer sets were used to generate the appropriate mutations within the gene fragment, which was cloned into the pRS404 vector ^20^. The vector was linearized and transformed into the desired yeast strain and colonies were selected on plates lacking tryptophan (the selectable marker in pRS404). After integration of the plasmid was confirmed by PCR, the strains were plated on media containing 5-fluoroanthranilic acid to select against vector sequence ^21^. Loss of vector was confirmed by PCR, and retention of the mutations was screened for by genomic PCR and confirmed by sequencing. For *in vivo* labeling of chromosomes, the plasmid pD215 targeted 256 *lac* operon operator (lacO) repeats ^22^ to the arm of chromosome *III* about 43 kbp from the centromere (coordinates 158504 -159234). Correct integration was confirmed genetically. A complete list of strains and their genotypes is shown in Table S1.

### Chromosome segregation assay

Strains were grown in YPAD medium at 30°C, except HJC2, DJC5, HJC14 and HJC22. These four strains were grown in a medium to select for cells that carried the mini-chromosome used to assay mis-segregation. Cells were diluted and plated on synthetic complete medium supplemented with 6 μg/ml adenine ^23 24^. This level of adenine allows growth of the *ade2-101* yeast cells that have lost the *SUP11* gene marker and permits accumulation of the red pigment used to identify segregation errors. Plates were incubated at 30°C until colonies were large (3-4 days) and further stored at 4°C for a few days to enhance differentiation of the color phenotypes. The half-sectoring assay was scored as previously described ^23^.

### Sister chromatid separation assay

Cells with P_*MET*_-*CDC20* (gift from Frank Uhlmann) ^25^ were arrested with 20 μg/ml α-factor in medium without methionine for 3 hr at 30°C. The cells were then washed with water and resuspended in medium without methionine. Methionine was added at 20 μg/ml as indicated to repress the P_*MET*_-*CDC20* promoter. Benomyl (#45339; Sigma-Aldrich) and nocodazole (#M1404; Sigma-Aldrich) were used 30 μg/ml of and 15 μg/ml, respectively, to depolymerize microtubules for 4 hr at 30°C. Nocodazole was re-added every hour at 7.5 μg/ml ^26^.

### Gamma-irradiation sensitivity test

Cells were grown in YPAD medium until mid-logarithmic phase, collected by brief centrifugation, washed with water, and resuspended in the original volume of water. The cells were exposed to a Cesium-137 source for the designated dosages. The average of three replicate platings were plotted. Each experiment was performed three times. One representative experiment is shown.

### Statistics and sequence analysis

Graphing and statistical analysis was done using Prism software and web-based tools (QuickCalcs), both from Graphpad (La Jolla, CA). Protein alignments were done with the ClustalW algorithm using MegAlign from DNAStar (Madison, WI). Parallel coiled-coil folds were predicted using Paircoil2 ^27^.

### Microscopy

Images were collected using a Carl Zeiss (Thornwood, NY) AxioImager microscope with band-pass emission filters, a Roper HQ2 charge coupled device, and AxioVision software.

## Results

### Some CdLS-associated mutations in *SMC1A* are defective for chromosome segregation

A relatively mild variant of Cornelia de Lange Syndrome (*CDLS2* [MIM: 300590]) has been attributed to mutations in the gene encoding the core cohesin subunit, Smc1 ^3,5,16^. Several of these mutations alter amino acid residues that are well conserved in Smc1 proteins from yeast to man, but their role in cohesin function is not well understood. It is not known whether any of these mutations affect the ability of the cohesin complex to tether sister chromatids together, or whether the phenotypes in affected individuals in fact reflect disruption of genome organization or transcriptional control rather than impaired cohesion *per se*.

One predictable outcome of reduced sister chromatid cohesion is increased chromosome mis-segregation and loss. Because these events in human cells are poorly tolerated and hard to score, we used budding yeast to better understand how CdLS-associated mutations in conserved amino acids of Smc1 affect cohesin function. To do this we used a sensitized assay that allows quantification of mis-segregation and loss of a mini-chromosome (CFIII, a truncated derivative of chromosome *III*) that is vulnerable, because of its small size, to defects in chromosome segregation machinery (Fig. 1) ^24^. This assay provides a colorimetric read-out for segregation of the marker mini-chromosome. Because the mini-chromosome carries no essential genes, cells that lose the mini-chromosome continue to propagate and can be scored in the mis-segregation assay. In diploid cells, the system allows the detection of both mis-segregation, in which two copies of the chromosome segregate to one daughter cell and none segregate to the other (2:0 segregation), and chromosome loss, in which one copy of the marker chromosome is lost during cell division (1:0 segregation) (Fig. 1). The mechanistic basis for chromosome loss is not clear, but it may reflect the failure of a chromatid, which has lost its association with its sister, to attach to microtubules from either side of the spindle, and to be left in the spindle mid-zone at anaphase I, thus failing to be included in either daughter nucleus ^23^. Both types of errors occur when cohesin fails. Thus, this system provides a sensitized assay for failures in establishing or maintaining sister chromatid cohesion.

**Figure 1.**
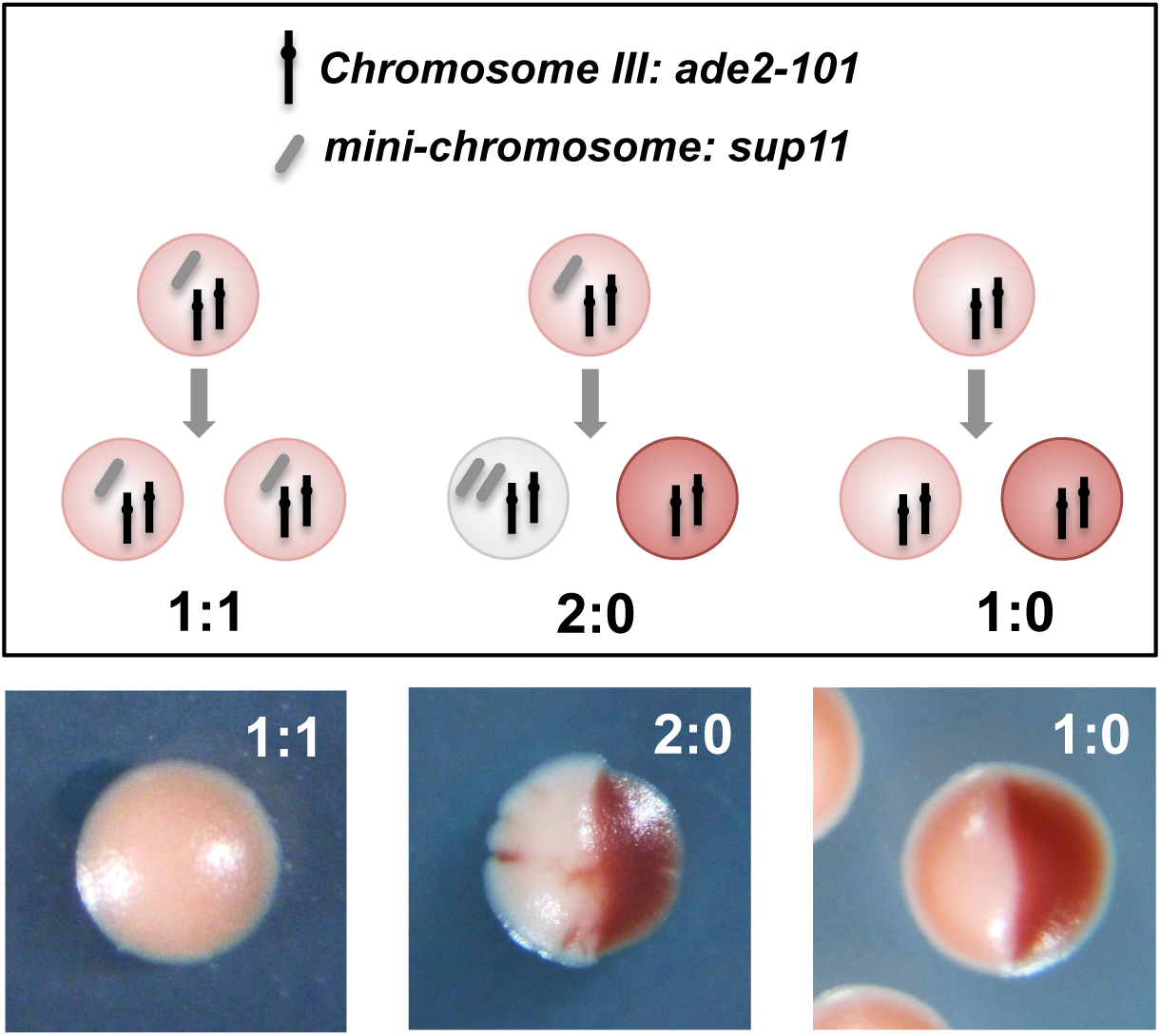
Illustration of the mini-chromosome segregation assay used to analyze cohesin function. **A.**Basis for the colorimetric readout ^23^. A marker mini-chromosome encodes a suppressor tRNA (*SUP11*), which suppresses the ochre nonsense mutation in an *ade2-101* allele carried on natural chromosome *III*. In the absence of suppression, *ade2-101* cells accumulate a pigmented intermediate in the adenine biosynthetic pathway, resulting in red cells and colonies. The copy number ratio between the marker mini-chromosome and the *ade2-101* allele determines the degree of suppression, and thus the colony color. Full suppression (1:1 ratio mini-chromosome:Chr *III*) results in white colonies, while partial suppression (1:2 ratio, mini-chromosome:Chr *III*) results in pink colonies, and in the absence of suppression (no mini-chromosome) colonies are red (Fig. 1). Colonies are scored based on the first segregation event (half colony only), and smaller sectors that arise later during outgrowth are ignored. Examples of sectored colonies are shown below the cartoon.

We chose for analysis ten mutations in *SMC1A* previously shown to lead to Cornelia de Lange Syndrome ^5^. These mutations map to both the coiled-coil region of the Smc1 protein, as well as to N and C terminal globular domains, and are associated with phenotypes of varying severity (Table 1) ^5^. Several of the affected residues are highly conserved throughout eukaryotic phylogeny, whereas some others are particularly well conserved in vertebrates (Fig. 2). To test the effects of these mutations on chromosome segregation, we generated analogous mutations in the yeast gene (Table 1), and integrated these mutations into the *SMC1* gene at its native locus, causing them to be expressed under control of the endogenous yeast *SMC1* promoter. We also generated a previously-characterized dominant-negative allele of *SMC1* to serve as a positive control for cohesin failure in the segregation assay. This mutation, *SMC1-657H* inhibits activity of a wild-type *SMC1* allele when co-expressed in the same cell, causing cohesion failure ^28^.

**Table 1.**
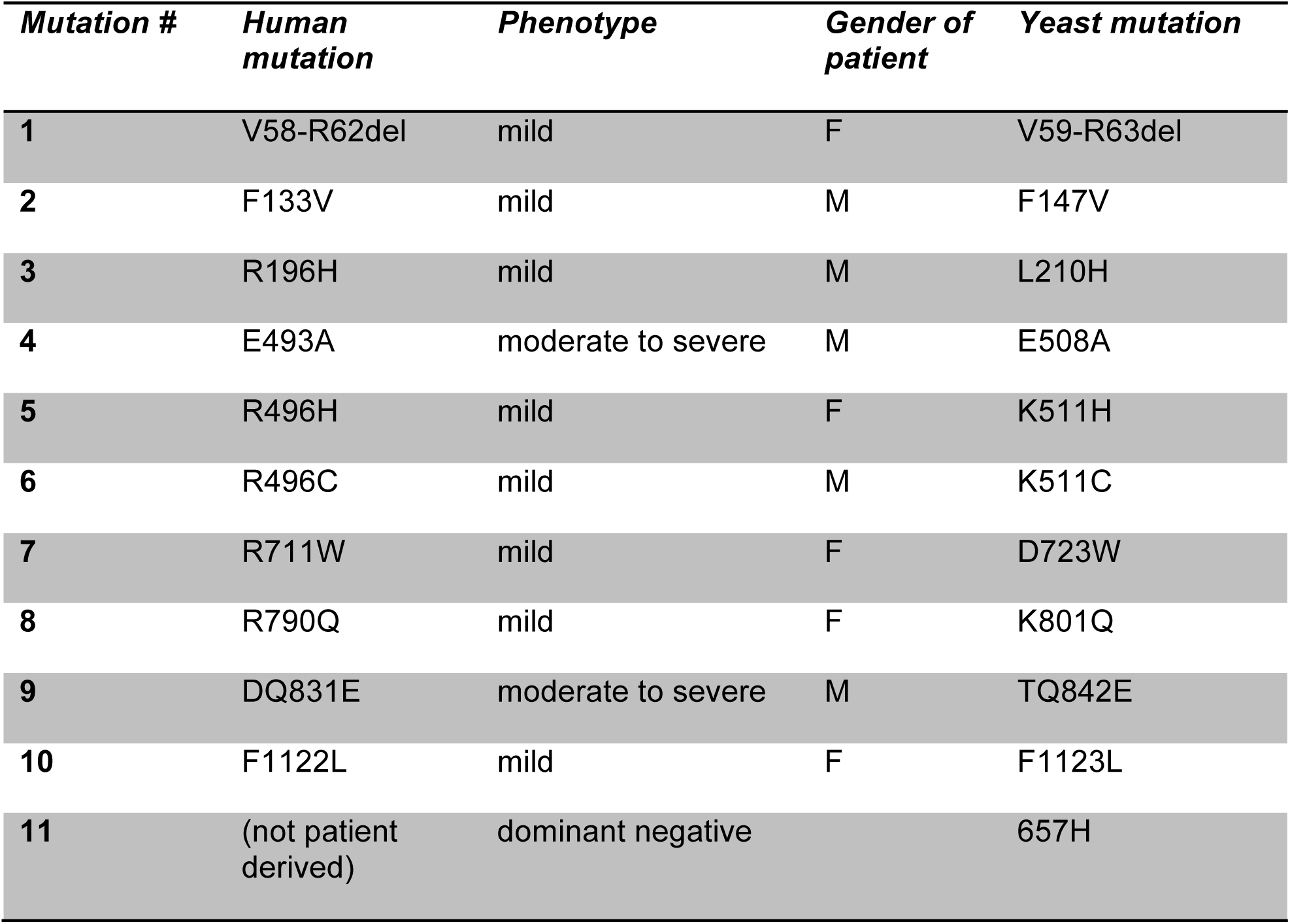
Mutations assessed in this study. Shown are the amino acid substitutions or deletions analyzed, and the severity of phenotypes in human patients as scored by Deardorff ^5^. The gender of the affected individuals in which the mutations were identified, and the analogous mutation in the yeast protein analyzed in this study, are indicated. The last mutation, 657H, is a dominant-negative allele of *SMC1* previously characterized in budding yeast ^28^.

| ***Mutation #*** | ***Human mutation*** | ***Phenotype*** | ***Gender of patient*** | ***Yeast mutation*** |
| --- | --- | --- | --- | --- |
| **1** | V58-R62del | mild | F | V59-R63del |
| **2** | F133V | mild | M | F147V |
| **3** | R196H | mild | M | L210H |
| **4** | E493A | moderate to severe | M | E508A |
| **5** | R496H | mild | F | K511H |
| **6** | R496C | mild | M | K511C |
| **7** | R711W | mild | F | D723W |
| **8** | R790Q | mild | F | K801Q |
| **9** | DQ831E | moderate to severe | M | TQ842E |
| **10** | F1122L | mild | F | F1123L |
| **11** | (not patient derived) | dominant negative |  | 657H |

**Figure 2.**
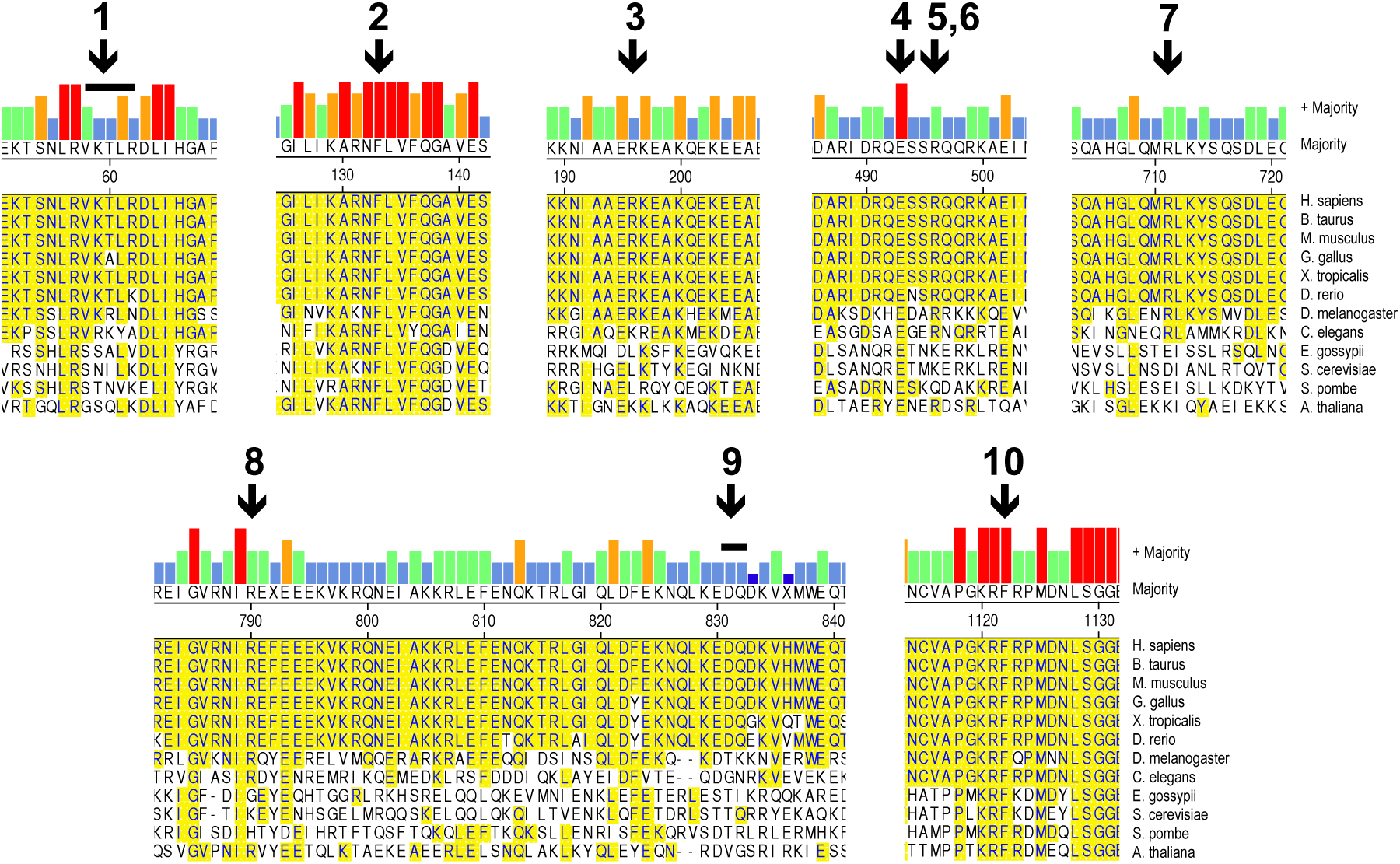
*SMC1A* mutations analyzed in this study. Shown is the sequence context of CdLS mutations in the human Smc1 protein. The positions of residues mutated in the human protein are indicated with black arrows labeled with bold font numbers. Small numbers indicate the sequence position in the human protein. Mutations that include deletions are indicated with black bars above the amino acids deleted. The mutation labeled **9** has both a single amino acid deletion and a substitution (*SMC1A-DQ831E*). ClustalW alignments of the relevant sequences in the human protein with Smc1 proteins from diverse eukaryotic organisms are shown. The strength of the alignment is indicated by the height of the colored bars at the top. Residues that match exactly the human sequence are highlighted in yellow. Full protein alignments are shown in Figure S1.

Versions of the *SMC1* gene bearing the mutations shown in Table 1 were introduced into diploid cells to create heterozygous cells (with one wild-type and one mutated copy of the gene). To test whether any of the mutations were lethal we sporulated the diploid strains, dissected the tetrads, and monitored viability of the haploid spores (half of which carry the wild-type allele and half of which carry the mutated version). All of the *SMC1* alleles tested, except *SMC1-657H*, supported viability of haploid strains and all exhibited colony sizes similar to the wild-type gene demonstrating that the CdLS-associated alleles do not cause catastrophic failures in sister chromatid cohesion.

Because Cornelia de Lange Syndrome-associated *SMC1A* mutations probably occur in a heterozygous configuration in females and are present as the sole allele of *SMC1A* in males, we scored the chromosome mis-segregation phenotypes of the alleles in both heterozygous and homozygous configurations. A schematic of the mis-segregation assay is shown in Figure 1. When screening *SMC1* alleles in the heterozygous configuration we compared their behavior to both the wild-type strain (*SMC1/SMC1*) as well as a heterozygous null strain, in which one copy of *SMC1* was deleted from the diploid background (*SMC1/smc1Δ*). Chromosome segregation was disrupted by four of the CdLS-associated alleles in the heterozygous configuration (Fig. 3A and Table S2). Heterozygous strains carrying the *smc1-TQ842-3E*, *smc1-Δ59-63*, *smc1-L210H*, and *smc1-F1123L* alleles all showed elevated rates of mis-segregation compared to the *SMC1/SMC1* control strain (P<0.05 for all but *smc1-F1123L*; Table S2). The *SMC1/smc1Δ* control also showed significantly higher error rates than the wild-type control demonstrating that Smc1 is a limiting component of the cohesin complex for maintaining sister chromatid cohesion. Earlier work had shown that levels of Scc1/Mcd1 subunit of cohesin can be lowered several fold with no clear impacts on sister chromatid cohesion, although these investigators used different, and probably less sensitive, assays to score cohesion function ^29^.

**Figure 3.**
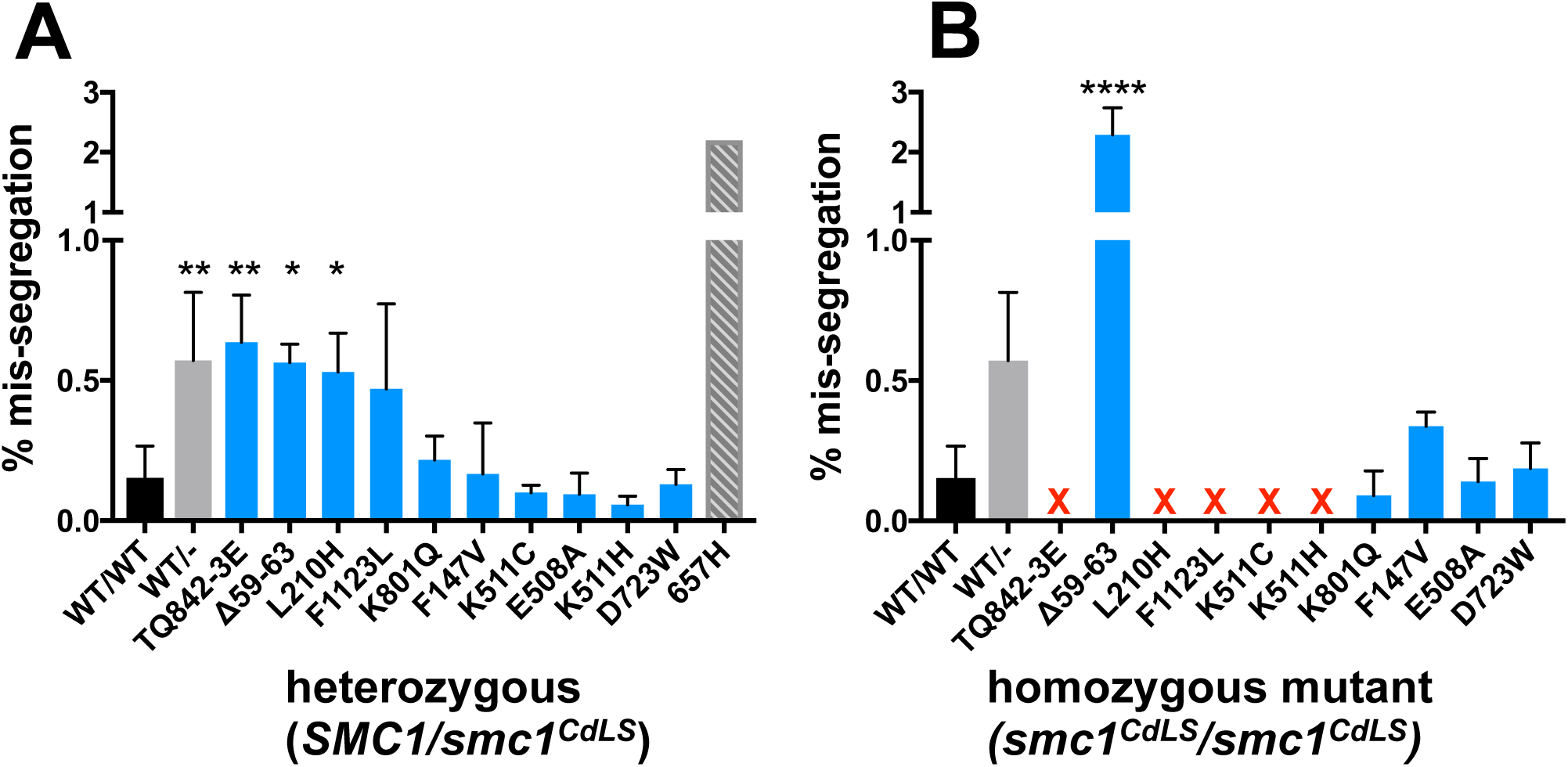
Chromosome mis-segregation in yeast strains carrying CdLS-associated mutations. **A.**Chromosome mis-segregation rates in *SMC1* WT/mutant heterozygotes. Shown are the percent of colonies exhibiting mis-segregation of the marker chromosome in the first cell division after plating. The total frequency of chromosome mis-segregation events (2:0 segregation and 1:0 segregation) are reported. Control strains are shown in black; solid black represents homozygous wild-type, while hatched black indicates heterozygous *SMC1/smc1Δ*. For a complete list of P values, see Table S2. **B.** Chromosome mis-segregation rates in mutant homozygotes (*smc1^CdLS^/ smc1CdLS*). Strains and controls are ordered as in **A.** For a complete list of P values, see Table S3. **C.** Chromosome mis-segregation rates in haploid mutant strains (*smc1^CdLS^*). Haploid mutant strains were similarly scored for chromosome mis-segregation events, and a haploid strain with wild-type *SMC1* was used as a control. In all graphs, chromosome mis-segregation rates that were statistically different from the *SMC1/SMC1* control are shown in red (Chi squared test with Yates correction, with Bonferroni’s correction for multiple comparisons). The previously characterized dominant negative mutant is shown in grey. **^*^** = strains that were not scorable due to extreme instability of the marker chromosome.

We found no evidence to suggest that any of the CdLS-associated alleles have dominant negative activity, as the chromosome mis-segregation rates were in no case worse than that of the *SMC1/smc1Δ* heterozygote. This indicates that the CdLS-associated alleles do not interfere (in a detectable way) with function of the wild-type protein present in the same cells. In contrast, and as anticipated, the known dominant-negative allele *SMC1-657H*, resulted in chromosome mis-segregation rates that were higher than that seen in the heterozygous *SMC1/smc1*Δ strain, consistent with previous conclusions that this mutation confers dominant-negative activity to the protein ^28^.

Six of the CdLS-associated mutations had no measurable effect on chromosome segregation when present with the wild-type allele as a heterozygote. These mutations could still compromise sister chromatid cohesion, but might be recessive to the wild-type allele, perhaps supplying sufficient partial function such that in the presence of a wild-type copy the cells have sufficient cohesion to behave like wild-type cells in the mis-segregation assay. To test this, we attempted to re-screen all of the alleles in the homozygous configuration in diploids (Fig. 3 B and Table S3). For several of the homozygous mutants this proved impossible, because the marker chromosome was lost so frequently that it was not possible to score the sectoring phenotype (Fig. 3 B, alleles marked by a red “X”). The four alleles that exhibited clear defects as heterozygotes (*smc1-TQ842-3E*, *smc1-Δ59-63*, *smc1-L210H*, and *smc1-F1123L*) also exhibited severe defects as homozygotes. In addition, strains bearing mutations at amino acid position 511, which had no clear defects as heterozygotes, exhibited severe chromosome mis-segregation defects as homozygotes. In contrast, strains that were homozygous mutant for *smc1-K801Q*, *smc1-F147V*, *smc1-E508A*, and *smc1-D723W*, showed chromosome mis-segrgation rates that were not different from the wild-type strain (Fig. 3B). This result is consistent with the lack of apparent mis-segregation defects seen when these mutations were present in the heterozygous condition (Fig. 3A).

### CdLS mutations in *SMC1* that cause mis-segregation have cohesion defects

Six of the CdLS alleles we assayed show very high levels of mis-segregation of the mini-chromosome in the assays shown in the sensitized system shown in Figure 3. The fact that cells carrying the *SMC1* CdLS alleles qualitatively exhibit near normal growth demonstrates that the mutations do not exert catastrophic defects on the behavior of natural chromosomes. To confirm that the mutant defects observed with the sensitized system reflect defects in sister chromatid cohesion, we performed a direct assay of natural chromosome behavior. We chose two of the mutations (*smc1-L210H* and *smc1-TQ842E*) that exhibited strong mis-segregation defects in the sensitized mini-chromosome segregation assay. We employed an assay that allows visual detection of cohesion failure, using a natural chromosome tagged on its arm with a fluorescent protein, GFP-LacI, that binds to an array of *lac* operon operator sequence repeats ^22,30^ (Fig. 4A). In cells with functional cohesion, the GFP foci on the cohered sisters appear as a single dot in most metaphase cells, whereas reductions in cohesin function allow the sister chromatids to separate, and two dots can be seen (Fig. 4 A) ^30^. To perform the assay, haploid cells were synchronized in G1 using alpha factor, released into the cell cycle and arrested in metaphase by limiting expression of *CDC20*, thereby preventing anaphase initiation. Metaphase cells were identified by the separation of the spindle pole bodies, marked by a red fluorescent protein, indicating that spindle had been formed. In the wild-type control (*SMC1*) only 2.81% of the chromosomes exhibited two GFP dots. In contrast both the *smc1-L210H* and *smc1-TQ842E* mutations resulted in greatly increased sister chromatid separation (45.2% and 64.4% respectively) (Fig. 4B). To test whether the sister chromatids were being pulled apart by the spindle or instead were perhaps never well tethered following their production during S-phase, the experiment was repeated in the presence of the microtubule poisons nocodazole and benomyl to disrupt spindle function. High levels of sister chromatid separation occurred even when spindle function was compromised by the addition of the microtubule poisons, demonstrating that the *smc1-L210H* and *smc1-TQ842E* mutations resulted in greatly impaired ability to ensure cohesion between sister chromatids of natural chromosomes.

**Figure 4.**
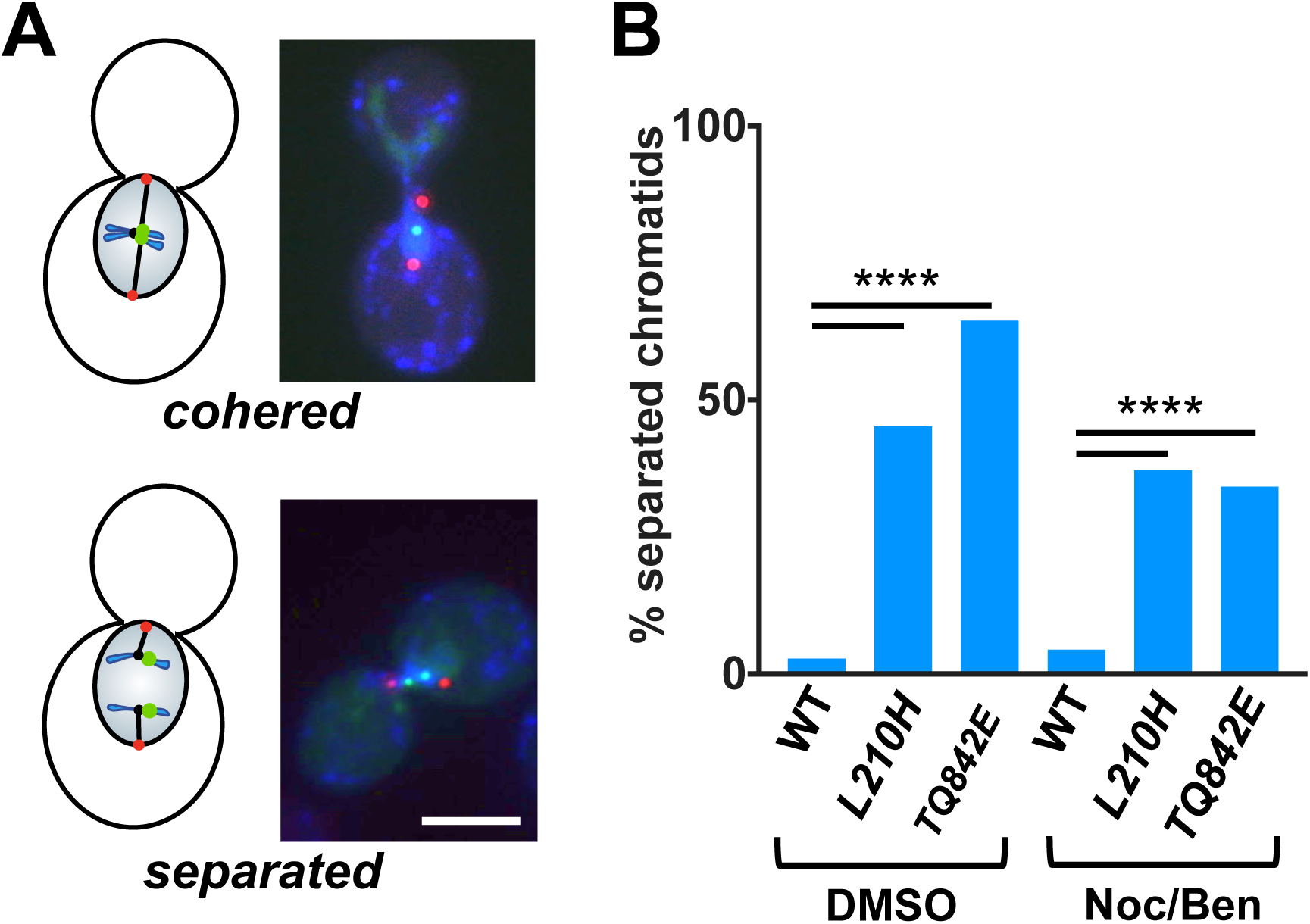
Precocious sister chromatid separation in yeast cells with CdLS mutations. **A.**Illustration of system used to monitor sister chromatid separation. Chromosome *III* harbors an integrated array of repeats of the *E. coli lac* operon operator sequence, which can be visualized as a green dot by expression of a protein fusion between green fluorescent protein and the lac repressor (GFP-lacI). The spindle pole body is tagged by expression of a fusion of the spindle pole body protein Spc42 with dsRed (Spc42-dsRed). In haploid cells, properly cohered sister chromatids appear as one green dot between two red dots in metaphase cells (top) while cohesion failure results in two green dots prior to anaphase (bottom). **B.** The level of sister chromatid separation in cells with different *SMC1* alleles. Haploid cells expressing the indicated alleles of *SMC1* were synchronized and allowed to progress into M phase in the presence of vehicle (DMSO, left) or nocodazole and benomyl (NocBen) for 4 hours, and the % of cells with separated green dots was measured. N>400 all samples. Fisher’s exact, ^****^ = 2 tailed P<0.0001.

### CdLS-associated mutations in *SMC1* increase sensitivity to ionizing radiation

Proper cohesion is essential for certain kinds of DNA repair, particularly those that involve homologous recombination ^31,32^. In budding yeast, DNA repair through homologous recombination induces *de novo* cohesion establishment throughout the nucleus ^31,33,34^. When cohesion establishment is compromised, survival is inhibited. To determine whether the CdLS-associated mutations in *SMC1* impact DNA repair, we exposed several strains bearing these mutations to ionizing radiation (IR) to generate DNA double strand breaks, and measured colony forming units as an indication of survival. We first tested heterozygous diploid *SMC1/smc1Δ* cells and compared them to the *SMC1/SMC1* (homozygous wild-type) cells to determine whether *SMC1* copy number impacts DNA damage sensitivity. In contrast to the mis-segregation assays, above, we found that the *SMC1* heterozygotes did not show increased radiation sensitivity compared to wild-type cells (Fig. 5A). This finding is in contrast to previous findings that DNA repair is more sensitive to reductions in Scc1/Mcd1 function than is chromosome segregation ^29^. This may reflect the differences in the specific assays used in the two studies or may suggest that the demands upon components of the cohesin complex may differ in mitotic segregation and DNA repair. Reasoning that the CdLS-associated mutations would therefore be unlikely to have a damage sensitivity phenotype in the presence of the wild type allele, we tested haploid cells bearing CdLS-associated mutations in the IR sensitivity. For this assay, we chose three of the CdLS mutations that had measurable impacts on chromosome segregation in the sectoring assay: *smc1-L210H*, *smc1-TQ842E*, and *smc1-F1123L* (Fig. 3). Consistent with a role for sister chromatid cohesion in DNA repair, all three of these mutations resulted in increased sensitivity to ionizing radiation (Fig. 5). The most severely affected strain carried the *smc1-TQ842E* mutation, which lost >90% viability following exposure to 600Gy of ionizing radiation. The *smc1-F1123L* and *smc1-L210H* mutations had intermediate phenotypes, losing ∼85% and 80% viability, respectively. In an earlier study, patient-derived cells bearing the human analog to the *smc1-F1123L* mutation were tested for IR sensitivity and found to be among the more IR sensitive of the small collection of *SMC1A* CdLS alleles tested, including the homologs of *smc1-K511H*, *smc1-K511C* and *smc1-V59-R63* ^35^. In that study, different cell lines bearing the same CdLS allele (*smc1-K511C* analog) showed different sensitivities suggesting that other genetic differences if the cell lines might influence the results. Here, because the strains are isogenic the observed differences are presumably solely due to the impacts of the different mutations. The IR sensitivity data presented here are consistent with the results in the sensitized chromosome segregation assay: the mutations tested impacted the ability of the cohesin complex to ensure accurate chromosome segregation and DNA repair and with the same order of severity. We conclude that these mutations impact both the ability of cohesin to tether sister chromatids together and to promote DNA repair.

**Figure 5.**
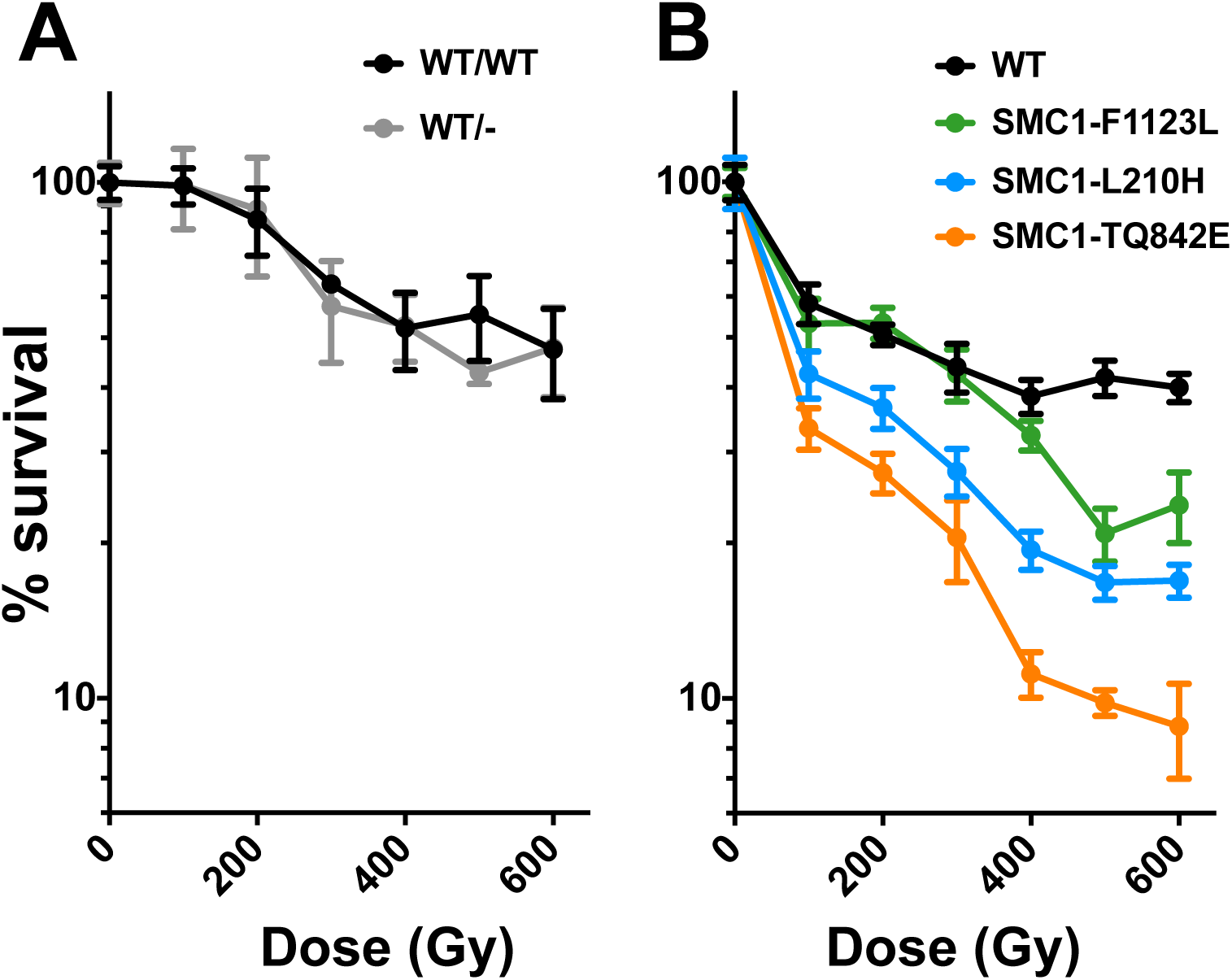
The effect of *SMC1* alleles on sensitivity to ionizing radiation (IR). **A.**Survival curves for diploid cells. Survival of diploid cells with the indicated genotypes following exposure to ionizing radiation was determined. **B.** Survival curves for haploid cells. Haploid cells of the indicated genotypes were exposed to ionizing radiation and survival was determined in a plating assay as in **A**.

## Discussion

The cohesion apparatus and its regulatory network are generally well conserved from yeast to man. We have analyzed the impact of a number of CdLS-associated mutations in human *SMC1A* on cohesin’s canonical functions by engineering analogous mutations into the budding yeast *SMC1* gene and measuring their impact on chromosome segregation and sensitivity to DNA damage. Our results suggest that these mutations can be broadly divided into two classes: those that have a clear measureable impact on chromosome segregation (and by extension sister chromatid cohesion) in budding yeast, and those that do not (Fig. 6). We interpret this to mean mutations of the second class impact functions of cohesin that are independent of its ability to tether together sister chromatids. Intriguingly, we found that two of the four CdLS-associated mutations in *SMC1A* (*smc1A-E493A* and *smc1A-R771*) that map to different ends of the primary protein sequence are predicted to map to the same position on the folded protein (Fig. 6). This is a region that is well conserved in vertebrates but less so among other species with the exception of the particular amino acid that is mutated (Fig. 2). It is possible that this region of the Smc1 protein, just adjacent to the coiled-coil and near the hinge domain, impacts development by directly affecting gene expression. This might be through direct interaction between cohesin and other factors that impact chromosome structure. Such models have been proposed, although we do not yet have a detailed mechanistic understanding of these interactions. The mediator complex, a global transcriptional regulator, co-purifies with cohesin, and like cohesin, contributes to the regulation of certain genes by stabilization of chromosome loops ^36^. It is possible that the interaction between cohesin and mediator occurs through the specific region in the coiled-coil of Smc1 highlighted in our analysis. An alternative model is that higher-order cohesin-cohesin interactions contribute to proper gene expression and thus development, and that specific CdLS-associated mutations in *SMC1A* disrupt these interactions ^37-39^.

**Figure 6.**
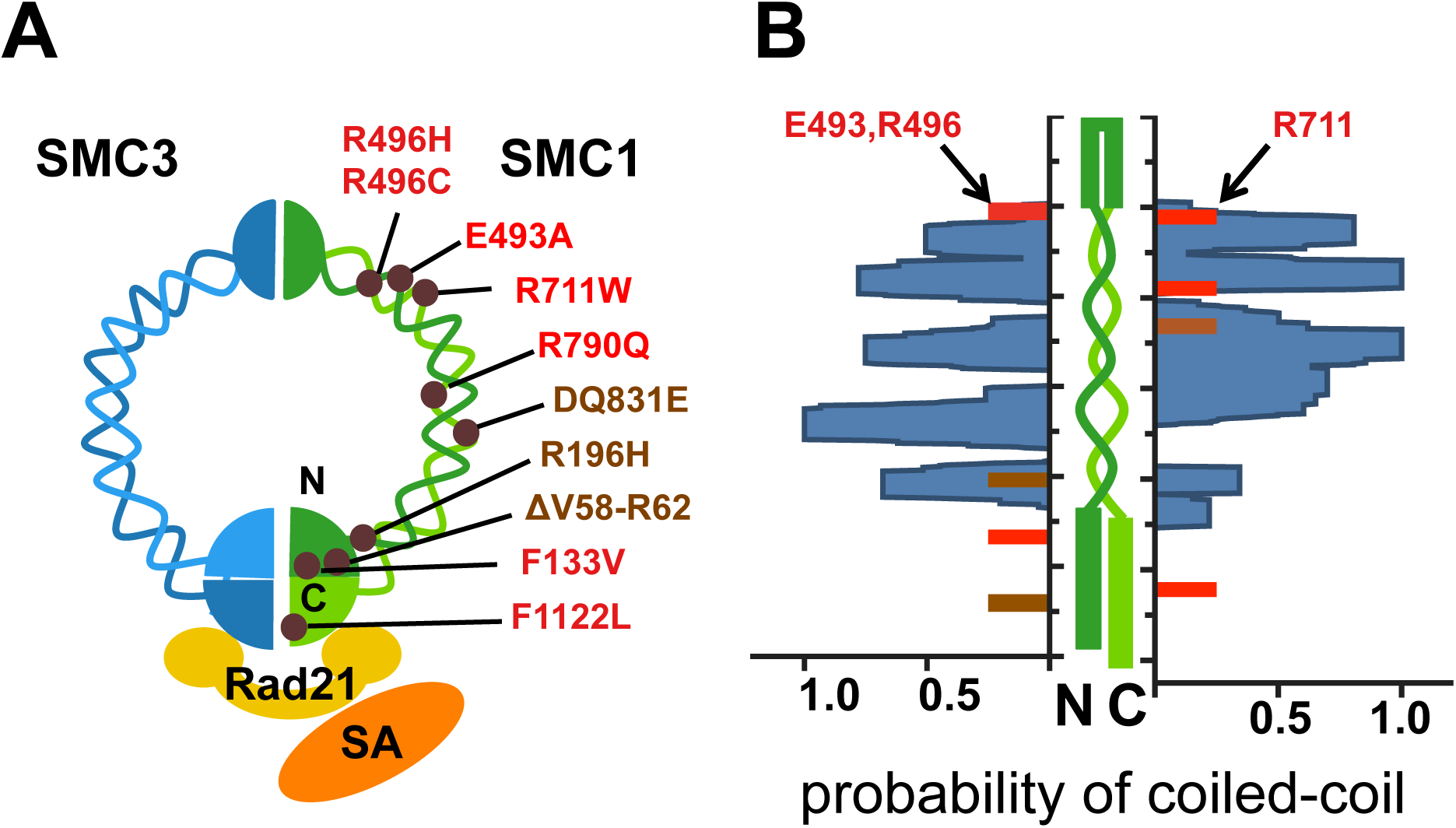
Cohesion-competent mutations map near each other in the folded *SMC1* protein. **A.**The approximate positions of individual CdLS-associated mutations tested in this study are shown in the context of the intact cohesin complex. Mutations with minimal effect on chromosome segregation in yeast-based assays are labeled in red; mutations that impact chromosome segregation are labeled in brown. **B.** Human Smc1 protein represented in green (center) with the N and C termini at the bottom and the hinge region at the top. The calculated probability of forming coiled-coil folds along the length of human Smc1 (Paircoil2; ^27^) is plotted along the length of human Smc1. The head and hinge domains of Smc1 are shown as green rods (bottom and top, respectively), while the coiled coils are shown as wavy green lines. Red bars indicate CdLS-associated mutations that have minimal impact on chromosome segregation in yeast assays. Brown bars indicate mutations that have defective chromosome segregation. Note that E493, and R711 align directly across from one another when coiled-coils are placed in register. Black ticks on vertical axes: 50 amino acid intervals. Probability scores (blue plot) are indicated at bottom.

We did not find that the severity of the CdLS phenotypes, as reported by Deardorff ^5^ correlated with the severity of the chromosome segregation defects in budding yeast. This supports the notion that cohesinopathies may reflect defects in cohesin functions that are distinct from its role in sister chromatid tethering. For example, the *smc1-E508A* mutation (corresponding to *SMC1A-E493A* in the human protein), had no measurable effect on chromosome segregation in the chromosome mis-segregation assay, but was among the most severe among the mutants that we tested in terms of the developmental phenotype in human patients ^5^. Conversely, some of the mutations with the severe chromosome mis-segregation phenotypes in our assays were identified in patients with relatively mild developmental phenotypes. One such mutation, L210H (analogous to R196H in humans) was identified in a male patient, and thus was present as the only *SMC1A* allele, though the phenotype was mild. Two of these mutations (*smc1-Δ58-62*, *smc1-F1122L*) map near the ATP binding site which is critical for sister chromatid cohesion ^40,41^, suggesting that proper interactions of cohesin with ATP are critical both for accurate chromosome segregation, and to prevent the symptoms of CdLS.

The mutations analyzed here have been identified in more than one family, and since beginning this work several additional *SMC1* mutations have been identified in CdLS patients^42^. Our results demonstrate that budding yeast provides a valuable tool with which to categorize mutations in cohesion proteins that impact human development. In the case of *SMC1*, using budding yeast to analyze mutations has allowed us to identify a region of the *SMC1* protein that may be particularly important for cohesin’s role in human development and distinct from its function in sister chromatid cohesion.

## References cited

1. Krantz, I.D., Mccallum, J., Descipio, C., Kaur, M., Gillis, L.A., Yaeger, D., Jukofsky, L., Wasserman, N., Bottani, A., Morris, C.A., et al. (2004). Cornelia de Lange syndrome is caused by mutations in NIPBL, the human homolog of Drosophila melanogaster Nipped-B. Nat Genet 36, 631–635.

2. Tonkin, E.T., Wang, T.-J., Lisgo, S., Bamshad, M.J., and Strachan, T. (2004). NIPBL, encoding a homolog of fungal Scc2-type sister chromatid cohesion proteins and fly Nipped-B, is mutated in Cornelia de Lange syndrome. Nat Genet 36, 636–641.

3. Musio, A., Selicorni, A., Focarelli, M.L., Gervasini, C., Milani, D., Russo, S., Vezzoni, P., and Larizza, L. (2006). X-linked Cornelia de Lange syndrome owing to SMC1L1 mutations. Nat Genet 38, 528–530.

4. Deardorff, M.A., Bando, M., Nakato, R., Watrin, E., Itoh, T., Minamino, M., Saitoh, K., Komata, M., Katou, Y., Clark, D., et al. (2012). HDAC8 mutations in Cornelia de Lange syndrome affect the cohesin acetylation cycle. Nature 489, 313–317.

5. Deardorff, M.A., Kaur, M., Yaeger, D., Rampuria, A., Korolev, S., Pié, J., Gil-Rodríguez, C., Arnedo, M., Loeys, B., Kline, A.D., et al. (2007). Mutations in cohesin complex members SMC3 and SMC1A cause a mild variant of cornelia de Lange syndrome with predominant mental retardation. Am J Hum Genet 80, 485–494.

6. McNairn, A.J., and Gerton, J.L. (2008). Cohesinopathies: One ring, many obligations. Mutation Research/Fundamental and Molecular Mechanisms of Mutagenesis 647, 103–111.

7. Mehta, G.D., Rizvi, S.M.A., and Ghosh,S.K. (2012). Cohesin: a guardian of genome integrity. Biochim Biophys Acta 1823, 1324–1342.

8. Ball, A.R., Chen, Y.-Y., and Yokomori, K. (2014). Mechanisms of cohesin-mediated gene regulation and lessons learned from cohesinopathies. Biochim Biophys Acta 1839, 191–202.

9. Merkenschlager, M., and Nora, E.P. (2016). CTCF and Cohesin in Genome Folding and Transcriptional Gene Regulation. Annu Rev Genomics Hum Genet 17, 17–43.

10. Harris, B., Bose, T., Lee, K.K., Wang, F., Lu, S., Ross, R.T., Zhang, Y., French, S.L., Beyer, A.L., Slaughter, B.D., et al. (2013). Cohesion promotes nucleolar structure and function. Mol Biol Cell.

11. Tomkins, D., Hunter, A., and Roberts, M. (1979). Cytogenetic findings in Roberts-SC phocomelia syndrome(s). Am J Med Genet 4, 17–26.

12. Rollins, R.A., Korom, M., Aulner, N., Martens, A., and Dorsett, D. (2004). Drosophila nipped-B protein supports sister chromatid cohesion and opposes the stromalin/Scc3 cohesion factor to facilitate long-range activation of the cut gene. Mol Cell Biol 24, 3100–3111.

13. Vrouwe, M.G., Elghalbzouri-Maghrani, E., Meijers, M., Schouten, P., Godthelp, B.C., Bhuiyan, Z.A., Redeker, E.J., Mannens, M.M., Mullenders, L.H.F., Pastink, A., et al. (2007). Increased DNA damage sensitivity of Cornelia de Lange syndrome cells: evidence for impaired recombinational repair. Hum Mol Genet 16, 1478–1487.

14. Kaur, M., Descipio, C., Mccallum, J., Yaeger, D., Devoto, M., Jackson, L.G., Spinner, N.B., and Krantz, I.D. (2005). Precocious sister chromatid separation (PSCS) in Cornelia de Lange syndrome. Am J Med Genet 138A, 27–31.

15. Kawauchi, S., Calof, A.L., Santos, R., Lopez-Burks, M.E., Young, C.M., Hoang, M.P., Chua, A., Lao, T., Lechner, M.S., Daniel, J.A., et al. (2009). Multiple organ system defects and transcriptional dysregulation in the Nipbl(+/-) mouse, a model of Cornelia de Lange Syndrome. PLoS Genet 5, e1000650.

16. Borck, G., Zarhrate, M., Bonnefont, J.-P., Munnich, A., Cormier-Daire, V., and Colleaux, L. (2007). Incidence and clinical features of X-linked Cornelia de Lange syndrome due to SMC1L1 mutations. Hum Mutat 28, 205–206.

17. Brown, C.J., Miller, A.P., Carrel, L., Rupert, J.L., Davies, K.E., and Willard, H.F. (1995). The DXS423E gene in Xp11.21 escapes X chromosome inactivation. Hum Mol Genet 4, 251–255.

18. Dresser, M.E., Ewing, D.J., Harwell, S.N., Coody, D., and Conrad, M.N. (1994). Nonhomologous synapsis and reduced crossing over in a heterozygous paracentric inversion in Saccharomyces cerevisiae. Genetics 138, 633–647.

19. Amberg, D.C., Burke, D., and Strathern, J.N. (2005). Methods in Yeast Genetics (CSHL Press).

20. Sikorski, R.S., and Hieter, P. (1989). A system of shuttle vectors and yeast host strains designed for efficient manipulation of DNA in Saccharomyces cerevisiae. Genetics 122, 19–27.

21. Toyn, J.H., Gunyuzlu, P.L., White, W.H., Thompson, L.A., and Hollis, G.F. (2000). A counterselection for the tryptophan pathway in yeast: 5-fluoroanthranilic acid resistance. Yeast 16, 553–560.

22. Straight, A., Belmont, A., Robinett, C., and Murray, A. (1996). GFP tagging of budding yeast chromosomes reveals that protein-protein interactions can mediate sister chromatid cohesion. Curr Biol 6, 1599–1608.

23. Hieter, P., Mann, C., Snyder, M., and Davis, R.W. (1985). Mitotic stability of yeast chromosomes: a colony color assay that measures nondisjunction and chromosome loss. Cell 40, 381–392.

24. Hegemann, J.H., Shero, J.H., Cottarel, G., Philippsen, P., and Hieter, P. (1988). Mutational analysis of centromere DNA from chromosome VI of Saccharomyces cerevisiae. Mol Cell Biol 8, 2523–2535.

25. Sullivan, M., and Uhlmann, F. (2003). A non-proteolytic function of separase links the onset of anaphase to mitotic exit. Nat Cell Biol 5, 249–254.

26. Fernius, J., and Marston, A.L. (2009). Establishment of cohesion at the pericentromere by the Ctf19 kinetochore subcomplex and the replication fork-associated factor, Csm3. PLoS Genet 5, e1000629.

27. McDonnell, A.V., Jiang, T., Keating, A.E., and Berger, B. (2006). Paircoil2: improved prediction of coiled coils from sequence. Bioinformatics 22, 356–358.

28. Milutinovich, M., Unal, E., Ward, C., Skibbens, R., and Koshland, D. (2007). A multi-step pathway for the establishment of sister chromatid cohesion. PLoS Genet 3, e12.

29. Heidinger-Pauli, J.M., Mert, O., Davenport, C., Guacci, V., and Koshland, D. (2010). Systematic reduction of cohesin differentially affects chromosome segregation, condensation, and DNA repair. Curr Biol 20, 957–963.

30. Michaelis, C., Ciosk, R., and Nasmyth, K. (1997). Cohesins: chromosomal proteins that prevent premature separation of sister chromatids. Cell 91, 35–45.

31. Unal, E., Arbel-Eden, A., Sattler, U., Shroff, R., Lichten, M., Haber, J.E., and Koshland, D. (2004). DNA damage response pathway uses histone modification to assemble a double-strand break-specific cohesin domain. Mol Cell 16, 991–1002.

32. Ström, L., Karlsson, C., Lindroos, H.B., Wedahl, S., Katou, Y., Shirahige, K., and Sjögren, C. (2007). Postreplicative formation of cohesion is required for repair and induced by a single DNA break. Science 317, 242–245.

33. Unal, E., Heidinger-Pauli, J.M., and Koshland, D. (2007). DNA Double-Strand Breaks Trigger Genome-Wide Sister-Chromatid Cohesion Through Eco1 (Ctf7). Science 317, 245–248.

34. Heidinger-Pauli, J.M., Unal, E., and Koshland, D. (2009). Distinct Targets of the Eco1 Acetyltransferase Modulate Cohesion in S Phase and in Response to DNA Damage. Mol Cell 34, 311–321.

35. Revenkova, E., Focarelli, M.L., Susani, L., Paulis, M., Bassi, M.T., Mannini, L., Frattini, A., Delia, D., Krantz, I., Vezzoni, P., et al. (2009). Cornelia de Lange syndrome mutations in SMC1A or SMC3 affect binding to DNA. Hum Mol Genet 18, 418–427.

36. Kagey, M.H., Newman, J.J., Bilodeau, S., Zhan, Y., Orlando, D.A., van Berkum, N.L., Ebmeier, C.C., Goossens, J., Rahl, P.B., Levine, S.S., et al. (2010). Mediator and cohesin connect gene expression and chromatin architecture. Nature 467, 430–435.

37. Huang, C.E., Milutinovich, M., and Koshland, D. (2005). Rings, bracelet or snaps: fashionable alternatives for Smc complexes. Philos Trans R Soc Lond B Biol Sci 360, 537–542.

38. Eng, T., Guacci, V., and Koshland, D. (2015). Interallelic complementation provides functional evidence for cohesin-cohesin interactions on DNA. Mol Biol Cell 26, 4224–4235.

39. Zhang, N., Kuznetsov, S., Sharan, S., Li, K., Rao, P., and Pati, D. (2008). A handcuff model for the cohesin complex. J Cell Biol 183, 1019–1031.

40. Weitzer, S., Lehane, C., and Uhlmann, F. (2003). A model for ATP hydrolysis-dependent binding of cohesin to DNA. Curr Biol 13, 1930–1940.

41. Arumugam, P., Gruber, S., Tanaka, K., Haering, C.H., Mechtler, K., and Nasmyth, K. (2003). ATP hydrolysis is required for cohesin′s association with chromosomes. Curr Biol 13, 1941–1953.

42. Gervasini, C., Russo, S., Cereda, A., Parenti, I., Masciadri, M., Azzollini, J., Melis, D., Aravena, T., Doray, B., Ferrarini, A., et al. (2013). Cornelia de Lange individuals with new and recurrent SMC1A mutations enhance delineation of mutation repertoire and phenotypic spectrum. Am J Med Genet 161A,2909–2919.

